# Robust adaptive distance functions for approximate Bayesian inference on outlier-corrupted data

**DOI:** 10.1101/2021.07.29.454327

**Authors:** Yannik Schälte, Emad Alamoudi, Jan Hasenauer

## Abstract

Approximate Bayesian Computation (ABC) is a likelihood-free parameter inference method for complex stochastic models in systems biology and other research areas. While conceptually simple, its practical performance relies on the ability to efficiently compare relevant features in simulated and observed data via distance functions. Complications can arise particularly from the presence of outliers in the data, which can severely impair the inference. Thus, robust methods are required that provide reliable estimates also from outlier-corrupted data.

We illustrate how established ABC distance functions are highly sensitive to outliers, and can in practice yield erroneous or highly uncertain parameter estimates and model predictions. We introduce self-tuned outlier-insensitive distance functions, based on a popular adaptive distance weighting concept, complemented by a simulation-based online outlier detection and downweighting routine. We evaluate and compare the presented methods on six test models covering different model types, problem features, and outlier scenarios. Our evaluation demonstrates substantial improvements on outlier-corrupted data, while giving at least comparable performance on outlier-free data.

The developed methods have been made available as part of the open-source Python package pyABC (https://github.com/icb-dcm/pyabc).

## 1 Introduction

Quantitative mathematical models are an indispensable tool in various research areas to describe and understand dynamical systems. Typically, models depend on unknown parameters that need to be inferred by calibration on experimentally observed data. The Bayesian paradigm allows to do so by combining the likelihood of observing data under given model parameters with prior information on the parameters, yielding a posterior distribution over parameters [Hines, 2015]. However, for complex stochastic models, evaluating the likelihood function is often infeasible [Jagiella et al., 2017, Tavaré et al., 1997, Wilkinson, 2009]. Approximate Bayesian Computation (ABC) is a likelihood-free method developed for such situations [Beaumont et al., 2002, Sisson et al., 2018, Tavaré et al., 1997]. In a nutshell, ABC compares summary statistics of simulated and observed data via a distance function and accepts if these are sufficiently close via some threshold, thus generating samples from an approximation to the posterior distribution. Due to its simplicity, scalability and broad applicability, it has become increasingly popular in various research areas, including e.g. population genetics, systems biology, ecology, epidemiology, and astronomy [Beaumont, 2010, Cameron and Pettitt, 2012, Imle et al., 2019, Sottoriva and Tavaré, 2010, Sunnåker et al., 2013, Syga et al., 2020]. ABC is frequently combined with a sequential Monte-Carlo scheme (ABC-SMC) [Del Moral et al., 2006, Sisson et al., 2007], which allows to efficiently iteratively reduce the acceptance threshold while maintaining high acceptance rates.

ABC relies on the ability to efficiently compare relevant features in simulated and observed data, via the combination of summary statistics and distance function. While both are in practice often still application-specific, various schemes have been devised to find appropriate low-dimensional summary statistics, e.g. via subset selection or regression [Blum et al., 2013, Fearnhead and Prangle, 2012]. As distance function, often a simple Minkowski distance is used, alternative approaches include e.g. Kullback-Leibler divergence [Jiang, 2018] or Wasserstein distances [Bernton et al., 2017] avoiding the use of summary statistics altogether. A popular distance function introduced by Prangle [2017] uses a weighted Euclidean distance that adjusts to the problem structure by iteratively updating summary statistic weights to normalize contributions, exploiting the structure of ABC-SMC algorithms.

The problem we tackle in this work is that errors can occur in the generation process of individual measurements, resulting in outliers in the collected data [Ghosh and Vogt, 2012, Motulsky and Christopoulos, 2003]. We informally denote by an outlier a data point corrupted by large errors that cannot be expected under the given experimental setup, already accounting for measurement noise. Reasons for outliers exist plenty, e.g. technical limitations, external perturbations, or human errors such as missing or incorrect labels [Maier et al., 2017].

Outliers can be problematic in parameter inference, as they can result in erroneous or highly uncertain parameter estimates. Consequently, methods have been developed that aim to detect and remove outliers, prior to and independent of the inference method used [Ben-Gal, 2005, Hodge and Austin, 2004, Niu et al., 2011]. However, for noise-corrupted high-dimensional or highly structured data with few replicates, which are common in biological problems and also as applications of ABC [Durso-Cain et al., 2021, Jagiella et al., 2017], such methods may be unreliable, and the complete removal of points that are not actually outliers can increase uncertainty [Motulsky and Christopoulos, 2003]. To circumvent this, estimators that are robust in the presence of outliers have been developed, using heavy-tailed distributions [Berger et al., 1994, Fernández and Steel, 1999, Huber et al., 1964, Tarantola, 2005] or pseudo-likelihoods with robust loss functions or divergences [Basu et al., 1998, Chérief-Abdellatif and Alquier, 2020, Jewson et al., 2018]. For ordinary differential equation (ODE) models of biochemical systems, the use of heavy-tailed likelihood functions such as Laplace, Huber, or Student’s *t*, rather than the commonly used normal distribution, has proven considerably more robust against outliers, without explicit removal [Maier et al., 2017].

Most such methods are however not applicable in ABC, as they require a tractable likelihood function. In this work, we propose a novel robust adaptive method for ABC, based on the adaptive distance function concept by Prangle [2017]. It can easily be shown that the original algorithm formulation is sensitive to outliers and can lead to biased or uncertain estimates (see Figure 1 for an illustration of possible problems on simple test models). We suggest the use of an outlier-insensitive norm, and additionally present a scheme for simulation-based active online outlier detection and correction.

**Figure 1:**
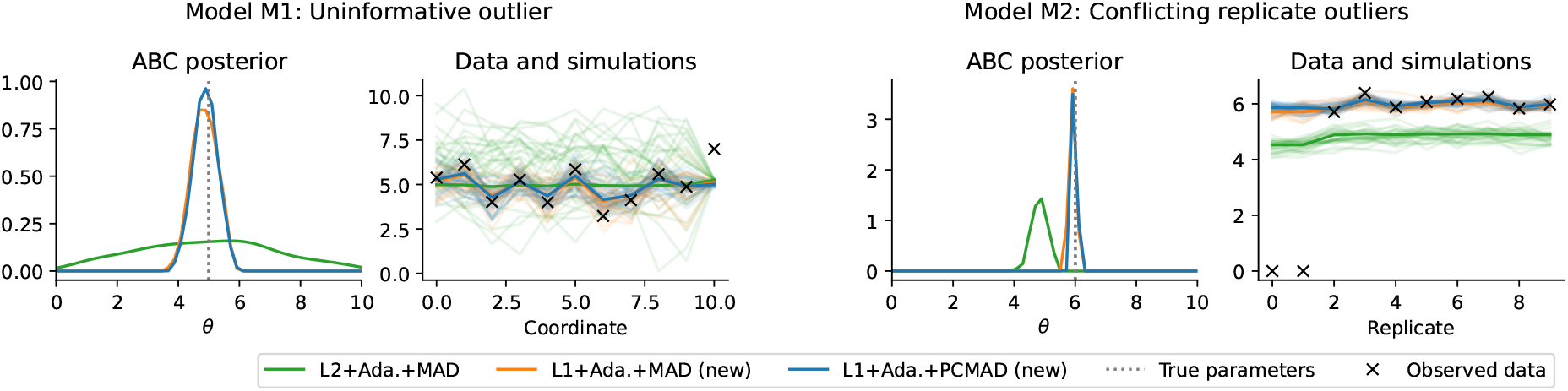
Problem illustration on two test models using three distances, the established L2+Ada.+MAD introduced in Prangle [2017], and two novel ones. Left: A model with 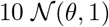 distributed data points, and an uninformative 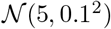 distributed one, whose observation however is an outlier. Right: A model with 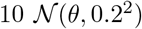 distributed data points, the first two of which however are outliers. For both test problems, on the respective left the obtained ABC posterior approximation is shown, and on the right the underlying outlier-corrupted data, together with, for all three distances, light-colored lines of 30 exemplary accepted simulations from the last ABC-SMC generation, and the respective sample means as darker lines. It can be seen how the established distance yields highly uncertain (left) or biased (right) estimates, while the novel methods give far more accurate estimates of the underlying true parameter. The distance functions and models shown here are properly introduced in Sections 2 and 3.

Existing robust inference approaches in ABC (not necessarily tailored to outliers) include notably Hellinger or Cramér-von-Mises distances as robust distances between probability distributions [Frazier, 2020], operating on high-dimensional data circumventing the definition of summary statistics, and approaches using distances based on *M*-estimating functions [Ruli et al., 2020] and *γ*-divergence estimators [Fujisawa et al., 2021]. The latter are however tailored to data with many i.i.d. replicates, which are not commonly available e.g. in systems biology applications. Another approach is presented in Frazier et al. [2020], who suggest to address model misspecification by parameterized adjustments of either the summary statistics or distance function weights. This approach is similar to ours in allowing the detection and down-weighting of inconsistent data points, and could in principle be combined with a scale adaptation scheme as done here. However, it augments the parameter vector by the number of summary statistics, thus appearing to require an appropriate, sufficiently low-dimensional, choice thereof. Lastly, a reformulation of the acceptance step as in Schälte and Hasenauer [2020] would allow for the use of heavy-tailed noise models similarly to Maier et al. [2017]. Yet, as in ABC the model must usually be regarded as a black box, such an approach is not generally applicable.

Our approach makes no assumptions on the type or dimension of data and is easy to adopt, allows for the interpretation of generated weights, and yields efficient inference by combination of robust measures with scale adaptation. This work focusing on the robust extension of the adaptive distance concept to outlier-corrupted data, a comparison to alternative ABC model misspecification approaches as mentioned above, in situations where several are applicable, would be of interest, but is beyond the scope of this work. We evaluate and compare the presented methods on various test examples covering various model types and features, and an application example of an agent-based model of tumor growth, on both outlier-free and outlier-corrupted data.

## 2 Methods

### 2.1 Background

#### 2.1.1 Approximate Bayesian Computation

Suppose we have measured data 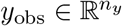, and have a descriptive model encompassing the underlying system dynamics, inherent stochasticity, and measurement noise. Bayesian inference combines the likelihood *π*(*y*|*θ*) of observing data 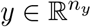 under the model, given model parameters 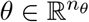, with prior information *π*(*θ*) on the parameters, to form a posterior distribution

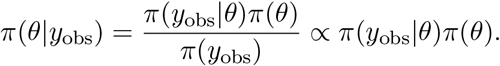

ABC deals with the situation that the model is generative, i.e. data *y* ∼ *π*(*y*|*θ*) can be simulated, but evaluating the (non-normalized) likelihood is infeasible. Classical ABC consists of the following three steps:

1. Sample parameters *θ* ∼ *π*(*θ*).
2. Simulate data *y* ∼ *π*(*y*|*θ*).
3. Accept (*θ, y*) if *d*(*s*(*y*), *s*(*y*_obs_)) ≤ *ε*.

Here, summary statistics 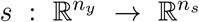 map the raw data to a typically lower-dimensional representation, 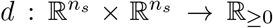 is a distance function measuring the “proximity” of simulated and observed data, and *ε* ≥ 0 is an acceptance threshold. This is repeated until sufficiently many, say *N* ∈ ℕ, particles have been accepted. Denoting *s* = *s*(*y*), *s*_obs_ = *s*(*y*_obs_), and 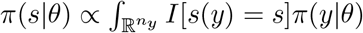 *dy* the intractable summary statistics likelihood, with *I* the indicator function, the population of accepted particles constitutes a sample from the approximate posterior distribution

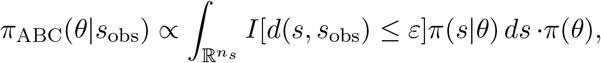

where 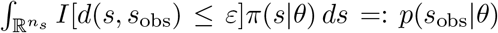 can be interpreted as an approximate likelihood function averaging over simulations proximate to *s*_obs_.

The uniform acceptance criterion *I*[*d*(·, *s*_obs_) ≤ *ε*] used throughout this work can be generalized to acceptance kernels *K*_*ε*_(*d*(·, *s*_obs_)) that do not only take a binary decision, but in the above third step accept with probability *K*_*ε*_(*d*(*s, s*_obs_))*/K*_*ε*,0_ [Sisson et al., 2018]. As we only deal with the choice of distance function, the methods presented in this work generalize to such kernels. If the ABC formulation does not employ an explicit distance function (such as Schälte and Hasenauer [2020]), the methods presented here do not apply. The methods presented here are moreover independent of the summary statistics method used.

#### 2.1.2 Sequential importance sampling

The above vanilla ABC algorithm, also called Rejection ABC, exhibits a trade-off between decreasing the acceptance threshold *ε* to obtain a better posterior approximation, and maintaining high-enough acceptance rates. To reconcile both, it is frequently combined with a Sequential Monte-Carlo (SMC) importance sampling scheme (Algorithm 1). In ABC-SMC, a series of particle populations 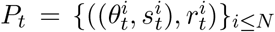 with acceptance thresholds 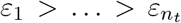, *t* = 1, … , *n*_*t*_, are generated, constituting samples from successively better posterior approximations. Particles for generation *t* are sampled from a proposal distribution *g*_*t*_(*θ*) ≫ *π*(*θ*) based on the previous generation’s accepted particles *P*_*t*−1_, e.g. via a kernel density estimate, only initially *g*_1_(*θ*) = *π*(*θ*). The importance weights 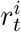 are the corresponding non-normalized Radon-Nikodym derivatives, *r*_*t*_(*θ*) = *π*(*θ*)*/g*_*t*_(*θ*).

An acceptance threshold scheme that has been shown to perform well and is also employed here, is to adaptively set *ε*_*t*_ to the quantile of the previous generation’s accepted distances 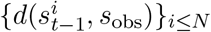 [Drovandi and Pettitt, 2011]. In addition, we automatically base the initial threshold on a calibration sample of size *N* that was also used to calibrate any other adaptive components, such as initial distance weights. A common form of the proposal distribution, which we also employed here, is 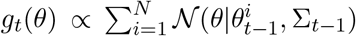 with covariance matrix 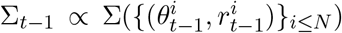 based on the sample covariance matrix. For details on the underlying ABC-SMC implementation used here see Klinger and Hasenauer [2017], Klinger et al. [2018].

For a test function 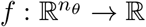, i.e. a statistic of interest, such as the mean or variance, the expected value under the approximate posterior is then approximated via the self-normalized importance estimator

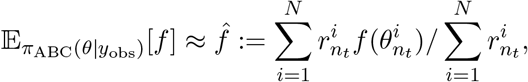

which is asymptotically unbiased as *N* → ∞ [Sisson et al., 2018].

##### Algorithm 1 ABC-SMC algorithm

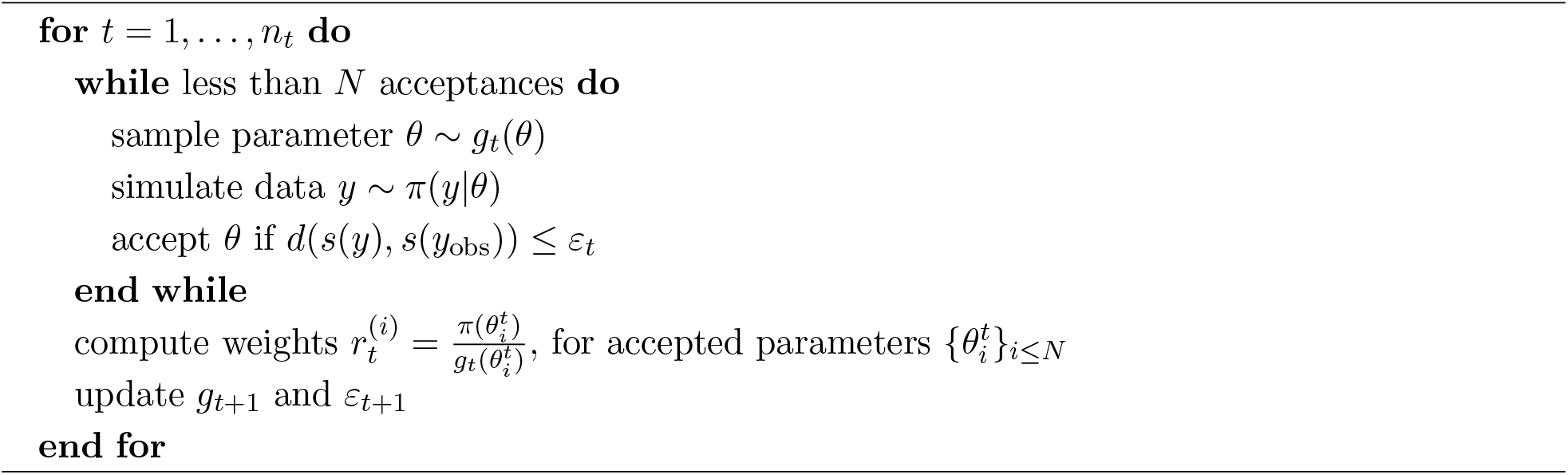

#### 2.1.3 Adaptive distances

A common choice of distance function *d* is a weighted Minkowski distance

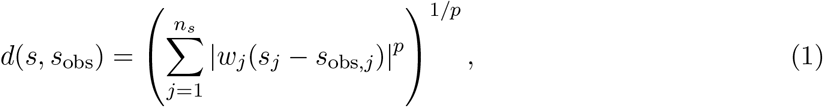

with *p* ≥ 1, where the summation is over the summary statistic coordinates *j*, and the *w*_*j*_ are summary statistic specific weights. It has been empirically argued that among similar distance functions, the exact form does not matter [McKinley et al., 2009, Owen et al., 2015]. Also theoretically, the approximate limit as *ε* → 0 is independent of the exact distance used (Section 2.2.3), if the data generation model is specified correctly [Schälte and Hasenauer, 2020].

However, as demonstrated in particular in Prangle [2017], adjusting the distance function to the problem structure can more efficiently yield substantially better parameter estimates, as more relevant information can be extracted from the data given a limited budget of simulations. The precise problem Prangle [2017] tackle is that summary statistics can vary on different scales. Thus, it can easily be that some statistics dominate the distance value, although the scale of a statistic is generally not informative of its relevance. This can be corrected for by the choice of the weights *w*_*j*_ in (1). A common choice is as inversely proportional to measures of variability of the respective statistics, i.e. *w*_*j*_ = 1*/σ*_*j*_ with *σ*_*j*_ e.g. given via empirical standard deviations 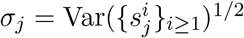, where {*s*^*i*^}_*i* ≥ 1_ are a calibration sample of summary statistics. An alternative measure suggested by Csilléry et al. [2012] for being more robust to sample outliers, and used in Prangle [2017] and also throughout this work, is the median absolute deviation (MAD) to the sample median,

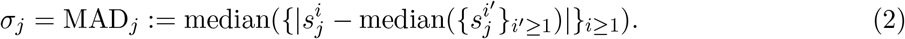

It is straightforward to base the weights on a calibration sample from the prior distribution, prior to the actual ABC analysis. However, while applicable to Rejection ABC, Prangle [2017] demonstrate that in an ABC-SMC framework, the distribution of summary statistics in later generations can differ considerably from prior samples. Their relative variability may change as the proposal distribution focuses on high-density regions in parameter space, such that weights obtained in calibration may no longer yield a similar contribution of each statistic to the distance value. Thus, they suggest the introduction of a sequentially updated generation-specific distance function *d*_*t*_, by updating the weights 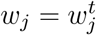 in distance (1) anew for each generation. The calibration sample for generation *t* consists of the particles generated in generation *t* − 1, including not only accepted ones but also rejected ones, in order to be representative of the sampling process and to allow greater flexibility of adaptation. This serves as an approximation to the expected summary statistic scale distribution under the proposal distribution *g*_*t*_ of generation *t*, without additional simulation costs. In order to ensure nested acceptance regions, Prangle [2017] then define the acceptance criterion for generation *t* as *d*_*t′*_ (*s, s*_obs_) ≤ *ε*_*t′*_ for all *t*′ ≤ *t*. When e.g. using an adaptive scheme for *ε*_*t*_, as mentioned in Section 2.1.2, it must be based on distance values recalculated with the new distance function, 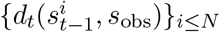.

While the above algorithm bases the weights for generation *t* on samples from generation *t* − 1 with a different proposal distribution, Prangle [2017] further propose a second algorithm that bases the weights on the current generation, by delaying the acceptance criterion definition. As this algorithm was found to not substantially improve results over the above introduced one, which is easier to integrate into existing ABC-SMC frameworks, we did not pursue this second formulation further.

### 2.2 Robust adaptive distances functions for outlier-corrupted data

We assume the data *y*_obs_ to be a realization of the model *π*(*y*|*θ*), except for single outliers that deviate from the actual trajectories and are due to another mechanism, as motivated in the introduction. In this section, we describe distance functions that build upon the adaptive distance functions introduced in Section 2.1.3, but are robust to outliers in the data, or can even correct for such.

#### 2.2.1 Outlier-insensitive absolute distances

While using MAD as a robust measure of sample variability for weight definition, Prangle [2017] base the overall distance on an L2 norm (*p* = 2 in (1)). Squared residuals emphasize large errors, which may be desirable, however makes the analysis highly sensitive to outliers, which typically result in large residuals that can dominate the distance and reduce the relative importance of other summary statistics, leading to an overall worse performance.

In regression analysis, robust approaches have been developed that are less sensitive to outliers than standard least squares. The arguably most common alternative are absolute deviations, corresponding to an L1 distance (*p* = 1 in (1)). For ODE models, it was shown in Maier et al. [2017] that replacing the most common assumption of a normal noise model, corresponding in a way to a weighted L2 distance, by heavy-tailed distributions such as Laplace, corresponding to an L1 distance, Huber, Cauchy or Student’s *t*, renders the analysis considerably more robust to outliers, while performing roughly comparably on outlier-free data. In ABC, if the noise assumption can be decoupled from the dynamics description, similar approaches can be employed, either by simulating e.g. Laplace measurement noise, or using an appropriate acceptance kernel [Schälte and Hasenauer, 2020]. However, in general, in ABC the model must be regarded as a black box that does not allow for decoupling of e.g. noise components. In that case, robustification and error correction must happen on the level of the distance function, which quantifies the impact of different data points.

Here, based on the above motivation, we propose to replace the L2 distance in the adaptive distance formulation by an L1 distance, yielding a distance function both robust to outliers and adaptive to the problem structure. In the following, we shortly motivate why this change of distance renders the analysis more robust to outliers. Consider the distribution of a single summary statistic given via a random variable *S* ∈ ℝ with variance 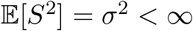, with observed value *s*_obs_. We are interested in the variability of the corresponding distance component in (1) around its average value, as similar levels of variability of different summary statistics yield similar impacts on the acceptance decision. For the variance of an L1 distance component |*S* − *s*_obs_| holds

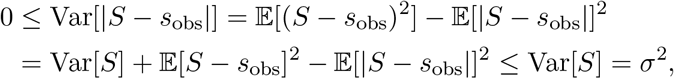

further Var[|*S* − *s*_obs_|] → *σ*^2^ in the limit |*s*_obs_| → ∞ of extreme outliers, with smaller variance for observed values closer to 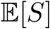. Thus, for simplicity assuming that in (1) normalization is by the distribution standard deviation, it follows for the variability of the corresponding component of the weighted L1 distance 0 ≤ Var[|*S* − *s*_obs_|*/σ*] ≤ 1. Note that the absolute value of the distance component may and will be larger for large outliers, however it only acts as an offset, while the variability is constrained. A similar boundedness does not generally apply to L2 distances. For example, assuming 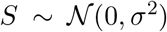, it is 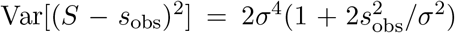, and thus for the weighted distance component 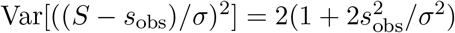, diverging for |*s*_obs_| → ∞.

#### 2.2.2 Online outlier detection and down-weighting via bias correction

Even when using an outlier-insensitive distance such as L1, large outliers can still impact the analysis, rendering it desirable to further detect and down-weight them. Here, we propose an online simulation-based approach to do so in ABC-SMC by complementing the MAD (2), as a measure of sample variability, by a measure of deviation from the observed value, such as the median absolute deviation to observation (MADO) for summary statistic *j*,

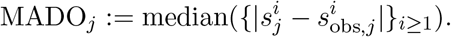

Accounting for both in-sample variability and deviation to observed value, we propose to then define the weight in (1) via the combined median absolute deviation (CMAD) as

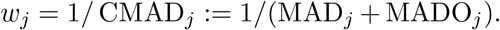

Outliers will typically have large deviation terms, which thus down-weight the statistic, such that the outliers impact the distance value and thus the analysis less. Conversely, if the model describes a data point well, one would expect the simulations to be close to the observed data, roughly on the same order as the in-sample variation of simulations, such that the additional term does not substantially hinder the purpose of variance normalization. Note that here we only down-weight data points while not removing them entirely as long as *w*_*j*_ ≠ 0, which conceptually permits the recovery from badly assigned weights at least in the approximate limit as *t* → ∞, i.e. no information is lost. Using an additive formulation instead of a multiplicative one, MAD_*j*_ · MADO_*j*_, ensures that the weights stay bounded in the presence of outliers even for MAD_*j*_ → 0.

Especially in early iterations and for uninformative priors, model simulations may deviate from the observed data substantially, as may the relative variability of different summary statistics. In order to not “punish” summary statistics that happen to deviate more from the observed data initially than others, we propose to apply the bias correction only if no more than a small fraction of the data points exhibit a substantial bias. Concretely, here we only used CMAD for weighting in a generation if *#*{*j* : MADO_*j*_ > 2 · MAD_*j*_}/*n*_*s*_ ≤ 1/3, resorting to MAD otherwise. These hyperparameters are clearly heuristics and may require tuning on some problems, however e.g. having more than one in three outliers in an analysis should in practice hardly occur, while only counting as outliers points with sufficiently high deviation compared to the in-sample variability focuses on large deviations that could otherwise considerably impact the analysis, while small outliers are less problematic (see also Section 3.7). We denote this weighting scheme, which perhaps uses CMAD and otherwise only MAD, as PCMAD.

As outliers usually get apparent merely in later generations, where simulations resemble the observed data more, applying such outlier correction methods only makes sense in an adaptive framework, as done here.

Instead of the robust measures MAD and MADO employed throughout this work, in principle also other measures of variability and deviation can be used, such as the common standard deviation and bias. We may thence statistically interpret the here introduced modifications as, informally, replacing the in-sample standard deviation by the root mean square error 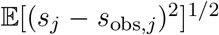, or robust alternatives thereof, treating the simulations as estimators of the observed value.

#### 2.2.3 Convergence

As noted in Prangle [2017] and also applicable here, convergence *π*_ABC_(*θ*|*s*_obs_) → *π*(*θ*|*s*_obs_) ∝ *π*(*s*_obs_|*θ*)*π*(*θ*) of the ABC-SMC posterior approximation as *t* → ∞ and *ε* → 0 is guaranteed if the adaptive distance metrics *d*_*t*_ are in particular of bounded eccentricity, i.e. the weight ratios bounded (note that generally only if *π*(*θ*|*s*_obs_) ≡ *π*(*θ*|*y*_obs_), i.e. the summary statistics are sufficient, is the original posterior recovered in the approximate limit). Practically, this can be achieved by constraining the relative weight range to a compact interval in (0, ∞), or e.g. by updating the distance function only a finite number of iterations. In the results presented in this work, we did not employ any such restrictions, as we focused on the ability of the methods to retrieve information under a limited budget of simulations. Yet, reliable strategies to set such constraints may be practically of relevance and may require further studies.

A further condition for convergence is naturally *π*(*s*_obs_) > 0, i.e. that the observed data can be simulated under the model, which may be impossible in the presence of outliers. The here-used down-weighting schemes however help to mitigate that problem, by focusing the analysis on reliable data points.

### 2.3 Implementation

The methods presented in this work have been implemented and made available in the open-source Python package pyABC (https://github.com/icb-dcm/pyabc), which implements a distributed ABC-SMC algorithm [Klinger et al., 2018]. Supplementary code of this study can be found at https://github.com/yannikschaelte/study abc rad, a snapshot of code and data is on ZEN-ODO at http://doi.org/10.5281/zenodo.5136475. All simulations were performed on the GCS Supercomputer JUWELS at Jülich Supercomputing Center (JSC), using up to 384 cores in parallel, with parallelization via dynamic scheduling [Klinger et al., 2018].

If not stated otherwise, like in Prangle [2017], we defined the acceptance criterion in generation *t* as *d*_*t′*_ (*s, s*_obs_) ≤ *ε*_*t′*_ for all *t*′ ≤ *t*, including previous acceptance criteria, to ensure nested acceptance regions. As transition kernel, we used a multivariate normal distribution with covariance kernel proportional to the previous generation’s weighted sample covariance matrix by Silverman’s rule of thumb. Acceptance thresholds were automatically selected as the median of the updated distance values 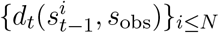 of the previous generation’s accepted particles. The distance weights were calculated based on all samples generated in the previous generation, including accepted and rejected ones.

## 3 Results

We tested the proposed methods on five test problems covering different problem features and model types, and a more realistic application problem, evaluating performance and robustness on both outlier-free and outlier-corrupted data, and comparing to established calibrated and adaptive distance functions as introduced in Prangle [2017].

### 3.1 Test models

An overview of the core problem features is given in Table 1. Details on all test problems can be found in the Supplementary Information, Section 1.

**Table 1:** Test model properties: Identifier, short description, number of parameters *n*_*θ*_ and data points or summary statistics *n*_*s*_, population size *N* and maximum number of model simulation after which an analysis was terminated.

| ID | Description | $n_\theta$ | $n_s$ | $N$ | Max. sim. |
| --- | --- | --- | --- | --- | --- |
| M1 | Informative and uninformative Gaussian variables | 1 | 11 | 1000 | 100000 |
| M2 | Independent Gaussian replicates | 1 | 10 | 1000 | 100000 |
| M3 | Conversion reaction ODE model | 2 | 10 | 1000 | 100000 |
| M4 | $g$-and-$k$ distribution order statistics | 4 | 7 | 1000 | 100000 |
| M5 | Lotka-Volterra Markov jump process model | 3 | 32 | 200 | 50000 |
| M6 | Tumor spheroid growth agent-based model | 7 | 150 | 500 | 150000 |

M1 consists of 10 observables 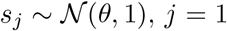, *j* = 1, … , 10 for a parameter *θ* with prior *θ* ∼ *U* [0, 10] and true value *θ* = 5, and one uninformative variable of low variance 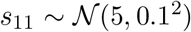. As outlier, we considered *s*_11_ = 7.

M2 consists of 10 observables 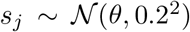, *j* = 1, … , 10, with prior *θ* ∼ *U* [0, 10] and true parameter *θ* = 6. While all observables are informative of the parameters, we set two to *s*_*j*_ = 0 to simulate conflicting information in the data due to outliers.

M3 is an ODE model of a conversion reaction 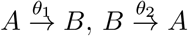, assuming species *B* to be observed at 10 evenly spaced time-points. We assumed additive noise 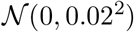. True parameters are (log *θ*_1_, log *θ*_2_) = (−1.5, −1.5), estimated on log-scale, with prior *U* [−3.5, 1]^⊗2^. Initial conditions were assumed to be known at (*A*_0_, *B*_0_) = (1, 0). As outlier, we randomly set one data point to zero, which could in practice e.g. occur as a wrongly assigned missing value. This model is identical to the first test model in Maier et al. [2017].

M4 and M5 are common ABC test problems, adopted directly from the application examples in Prangle [2017]. M4 is based on the g-and-k distribution, which is defined via its quantile function but does not have a closed-form likelihood function. We used as summary statistics seven order statistics at indices (1250, 2500, …, 8750) out of 10, 000 independent samples. We used independent *U* [0, 10] priors on the model’s four parameters *A, B, g, k*, and ground truth values (*A, B, g, k*) = (3, 1, 1.5, 0.5). As outlier we considered randomly setting one observable to zero. M5 is a Markov jump process model of a Lotka-Volterra predator-prey population model, simulated via Gillespie’s direct algorithm [Gillespie, 1977]. We assumed both species to be observed under addi-tive 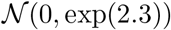 distributed noise, at 16 evenly spaced time-points over a span of roughly four periods. The model has three conversion rate parameters, which were estimated on log-scale, with wide independent *U* [−6, 2] priors and ground-truth values (*θ*_1_, *θ*_2_, *θ*_3_) = (1, 0.005, 0.6). As outliers we considered multiplying 6 observables at 3 random time-points by a factor of 10, which can in practice occur e.g. due to a wrong exponent.

M6, a more realistic application example, is a multi-scale agent-based model of tumor spheroid growth on a two-dimensional plane, as introduced in Jagiella et al. [2017]. Single cells are modeled as stochastically interacting agents, coupled to the dynamics of extracellular substances modeled via partial differential equations. The model describes three observables: The spheroid radius over time, and the extra-cellular matrix (ECM) density and the fraction of proliferating cells, at different distances from the rim, observed at a single time-point. On top of the agent-based model, we assumed independent normal measurement noise. The model possesses seven unknown parameters. As outliers we considered interchanging in total 20 data points in the observables’ dynamic regimes. Agent-based models describing biological processes such as pathogen spread or tissue growth have recently been frequently analyzed using ABC methods [Durso-Cain et al., 2021, Imle et al., 2019].

### 3.2 Experimental setup

To assess the performance of the various described distance functions, we ran the first five test models 20 times on outlier-free data, each for a separate data set randomly sampled from the model under ground truth parameters *θ*_GT_, and additionally 20 times on outlier-corrupted data sets derived from the outlier-free ones. Each run was performed with a fixed population size *N* and given a budget of model evaluations, evaluated after each generation, as given in Table 1. The evaluation budgets were chosen rather low, in order to assess the ability of the distance functions to extract information from simulations under a limited budget.

We evaluated the performance of a distance function by its ability to yield accurate point estimates with low uncertainties. Thus, we used, as a metric combining both aims, the root mean square error (RMSE) of the weighted posterior samples from the last ABC-SMC iteration with respect to the ground truth parameters, i.e. the square root of 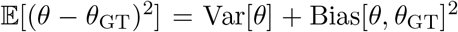 under the obtained posterior approximation. While the actual posterior mean may not be unbiased for a given data set, one may expect it to be on average.

As distance functions, we considered L2 and L1 distances (“L2” or “L1”), only pre-calibrated in the first iteration or adapted in each generation (“Cal.” or “Ada.”), and using MAD, CMAD, or PCMAD for distance weight calculation. In the following, e.g. “L2+Ada.+MAD” denotes an adaptive L2 distance with MAD-based weights. While we also explore other combinations, we focus the analysis on the established pre-calibrated L2+Cal.+MAD and adaptive L2+Ada.+MAD, and the here newly introduced robust L1+Ada.+MAD and L1+Ada.+PCMAD.

### 3.3 L1 comparable to L2 on outlier-free data, and clearly outperforming on outlier-corrupted data

Comparing the performance of L2- and L1-based distances (Figure 2, upper half vs. lower half) on outlier-free data (Figure 2, light bars), e.g. for L2+Ada.+MAD vs. L1+Ada.+MAD, shows no substantial differences for most models and parameters, indicating that L1 works similarly well on outlier-free data.

**Figure 2:**
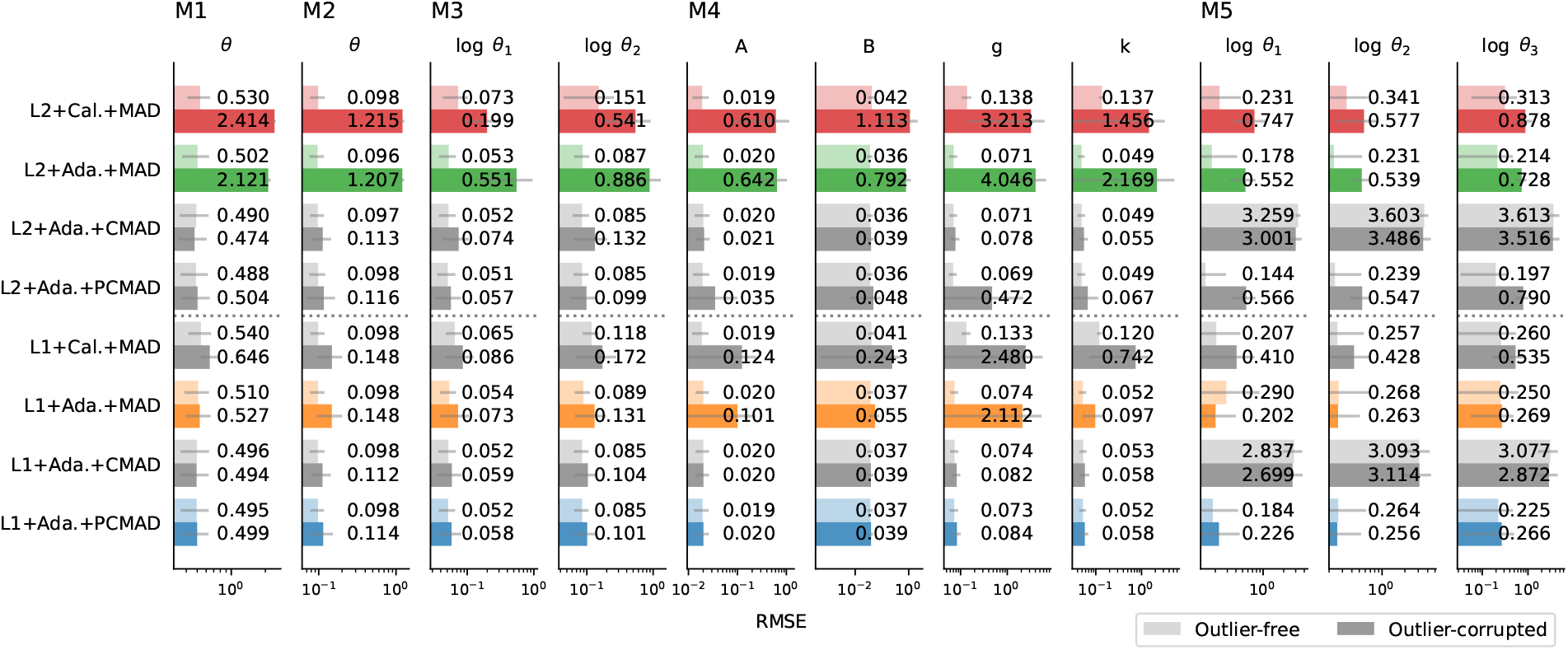
Mean RMSE for the parameters of 5 test models (columns) obtained for 8 distance functions (rows), using L2 or L1 distances, calibrated only in the first (“Cal.”) or every (“Ada.”) generation, and using MAD, CMAD, and PCMAD for distance weight calculation. Each RMSE is averaged over 20 data sets, grey lines indicate standard deviations. For each distance, the upper, lighter bar is based on outlier-free data, while the lower, darker bar is based on outlier-corrupted data. Distances of interest are colored, alternative distance combinations are shown in grey for reference.

On outlier-corrupted data (Figure 2, dark bars), using an L2 distance (L2+Ada.+MAD) yielded substantially higher RMSE values compared to outlier-free data. Using an L1 distance (L1+Ada.+MAD) drastically reduced the RMSE in all cases by up to orders of magnitude, in many cases to only slightly higher levels than obtained by the same distance for outlier-free data.

Comparing pre-calibrated L2+Cal.+MAD and adaptive L2+Ada.+MAD (as well as the L1 variants) on outlier-free data confirms the finding in Prangle [2017] that the adaptive weighting scheme can better adjust to the problem and thus outperforms pre-calibrated weighting. However, e.g. the comparison of L2+Cal.+MAD and L2+Ada.+MAD on outlier-corrupted data shows that there, in some cases, an adaptive weighting scheme with L2 distances can give worse results than pre-calibrated weights, arguably due to decreasing in-sample variability with large impact on the distance function, as outlined in Section 2.2.1. For the corresponding L1 distances, conversely in the majority of cases, an adaptive distance function improved results over pre-calibration.

### 3.4 Outlier correction further improves estimates and identifies outliers

Applying active outlier detection and down-weighting by PCMAD on top of using an L1 distance (L1+Ada.+PCMAD) generally further reduced RMSE values over MAD (L1+Ada.+MAD) on M1-4, and substantially so e.g. for the *A* and *g* parameters of problem M4, giving a total reduction of up to nearly 50 times compared to the established L2+Ada.+MAD. On M1-4, the use of CMAD throughout (L1+Ada.+CMAD) and not only if outliers were detected in less than one in three data points (PCMAD), yielded low RMSE values as well. However, the inverse was true for M5, where L1+Ada.+CMAD gave large RMSE values on both outlier-free and outlier-corrupted data. This is likely due to the fact that M5 exhibits highly flexible dynamics, with trajectories under the prior predictive distribution being considerably different from the posterior. In that situation, blindly applying bias correction may wrongly down-weight data points that just happen to not have converged yet, while PCMAD reliably only does so in case of strong indication of few outliers.

Overall, in many cases L1+Ada.+MAD already gave substantially better results than the reference method L2+Ada.+MAD, which could be further improved by outlier detection, such as here L1+Ada.+PCMAD, which consistently yielded small RMSE values for all parameters of all test problems.

### 3.5 Nested acceptance regions improve performance slightly

In order to assess the effect on robustness of nested acceptance regions via checking previous acceptance criteria, *d*_*t′*_ (*s, s*_obs_) ≤ *ε*_*t′*_, *t*′ ≤ *t*, and not only the current criterion, we ran the same analyses as shown here, but only taking into account the current generation’s acceptance criterion, i.e. accepting if *d*_*t*_(*s, s*_obs_) ≤ *ε*_*t*_ (Supplementary Information, Section 2). Overall, we found no major or structured differences between using previous acceptance criteria or not, especially for L1 distances, on both outlier-free and outlier-corrupted data, with slightly better results when using previous acceptance criteria. An exception, where the use of previous acceptance criteria performed substantially better, was M5, which especially for L2 showed a larger RMSE for the adaptive distance compared to the pre-calibrated distance when not doing so. A reason for this may be the highly flexible model dynamics under the wide prior, for which a more conservative acceptance criterion may be more robust. Both using and discarding previous acceptance criteria may have their advantages – using them may be more conservative and thus preferable on highly variable models, while not using them gives more flexibility and may allow to escape from bad initial choices.

### 3.6 Robust distances yield better fits of the data

Not only in terms of parameter estimates, but also in terms of fits of simulated data to the observed data, and thus ultimately in terms of predictions, did the advantage of the robust adaptive distance functions L1+Ada.+MAD and L1+Ada.+PCMAD over L2+Ada.+MAD become apparent (shown in Figure 3 exemplarily for selected data sets). For M1, the variability of simulated data for L2+Ada.+MAD was considerably higher than for the other distances. For M2, L2+Ada.+MAD tried to balance between the eight accurate measurements and the two outliers, fitting no point well, while L1+Ada.+MAD showed considerably less bias, only slightly more than L1+Ada.+PCMAD. This held similarly for M3 and M4, where the analysis for L2+Ada.+MAD converged to curves visibly differently shaped than the observed data, corresponding to different parameter vectors. For M5, extremes of the dynamics were less well captured. Overall, L1+Ada.+PCMAD gave the best fits in all these cases.

**Figure 3:**
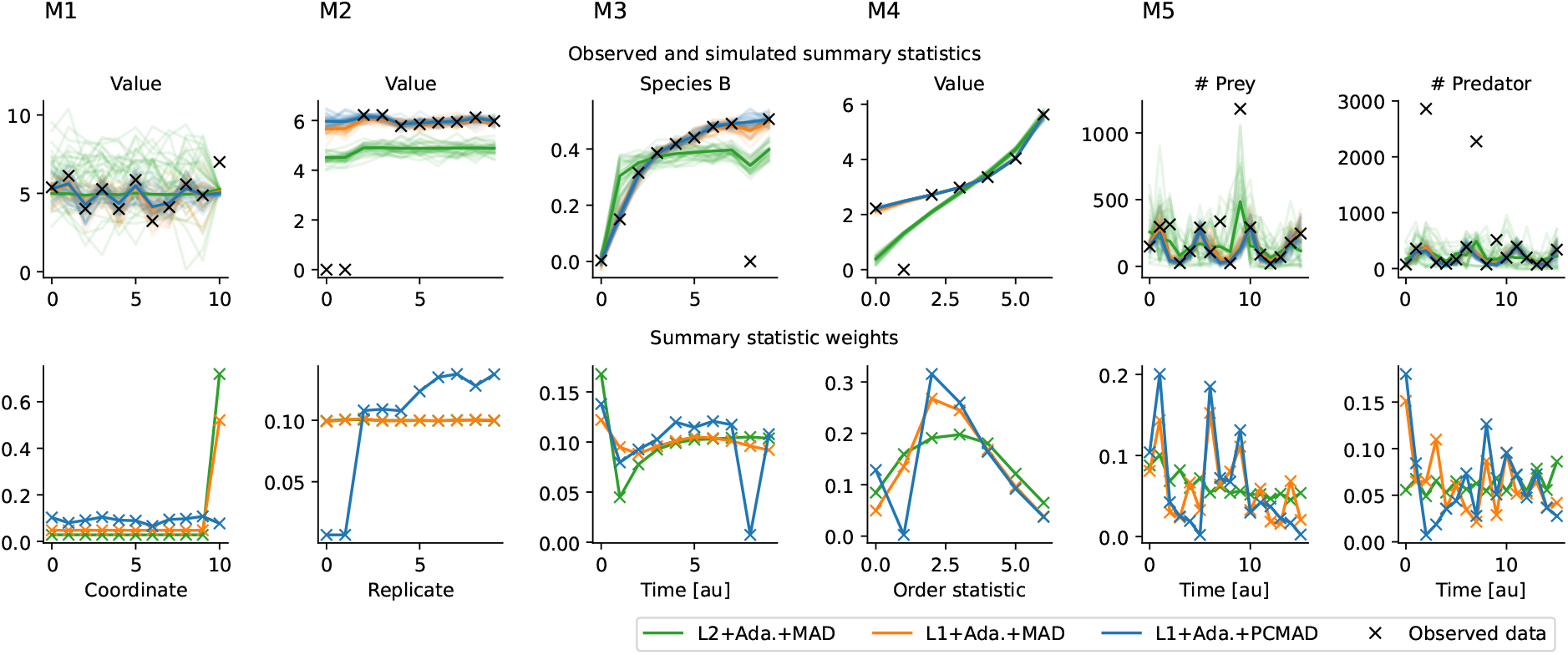
Fits and weights for models M1-5 on outlier-corrupted data, for three distances. Upper row: Observed data (black) and, for each distance, 30 accepted simulated data sets (lighter lines) as well as the sample means (darker lines) from the last ABC-SMC generation. Note that these are accepted simulations, not predictions; for *ε* → 0, the accepted simulations should exactly match the observed (non-outlier) data. Lower row: The corresponding weights assigned to each summary statistic by the three shown distance functions in the last generation, normalized to sum 1. For each problem, one exemplary run out of the 20 runs on outlier-corrupted data is shown.

This was also reflected in the weights the respective distances assigned to data points in the last ABC-SMC generation (Figure 3 bottom). For M1-4, PCMAD reliably detected outliers and assigned them low weights, decreasing their impact on the analysis. Typically, bias correction was applied only after a few iterations due to overall high initial variability, reliably converging to the actual outliers in the later iterations in most cases. For M5, the PCMAD automatic outlier correction did not generally identify all outliers. This is in line with the previous finding that here PCMAD does not give an advantage over MAD alone, while blindly applying CMAD is detrimental. Likely, the flexibility of the model does not allow to reliably detect outliers, as too many data points exhibit deviations from the observed values.

### 3.7 Robust to outliers at large scale

To analyze the impact of the scale of outliers on the presented methods, we ran the test problems M1 and M2 with outliers at different multiples of the corresponding statistics’ standard deviations (Supplementary Information, Section 3). This revealed that the performance of L2+Ada.+MAD considerably worsened with outliers at larger scales, while the L1-based distances gave substantially more reliable results, with slight improvements for PCMAD over MAD. This is in line with the argumentation of Section 2.2.1.

### 3.8 Robust methods clearly outperform established methods on complex application example

Due to its computational complexity with model simulation times on the order of seconds, for the tumor growth application problem M6 we only used single outlier-free and outlier-corrupted data sets. Here, we compared a pre-calibrated distance L2+Cal.+MAD, an adaptive L2 distance L2+Ada.+MAD, and the newly introduced L1 distances L1+Ada.+MAD and L1+Ada.+PCMAD.

The parameter estimates show that, remarkably, the L1 distances performed superior to L2 on M6 also on outlier-free data (Figure 4 top). In particular L2+Ada.+MAD yielded substantially wider uncertainty estimates, also compared to L2+Cal.+MAD. This is also reflected by the model simulations being more dispersed (Figure 5 top). Arguably, this is due to the fact that for high-dimensional data the chance of deviant data points is high, even if the model perfectly describes the underlying data generation process. In that case, an outlier-sensitive adaptive L2 distance may increasingly focus the analysis on points with large deviation, biasing the analysis, even if there are no systematic outliers in the data.

**Figure 4:**
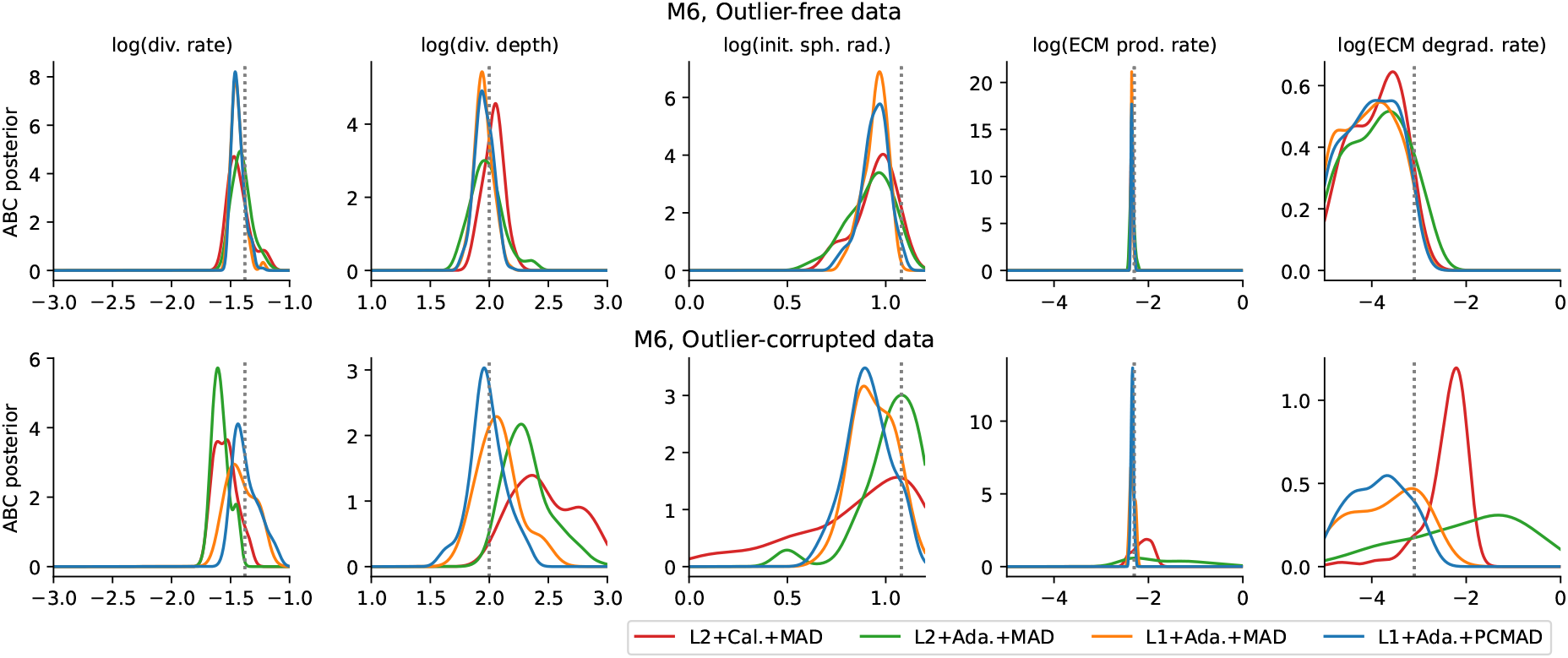
Posterior marginals for the 5 out of the 7 model parameters of model M6 showing interesting dynamics. Top row: Without outliers. Bottom row: With outliers. The x-axis boundaries are the uniform prior boundaries. The parameter values used to simulate the observed data are indicated by grey dotted lines.

**Figure 5:**
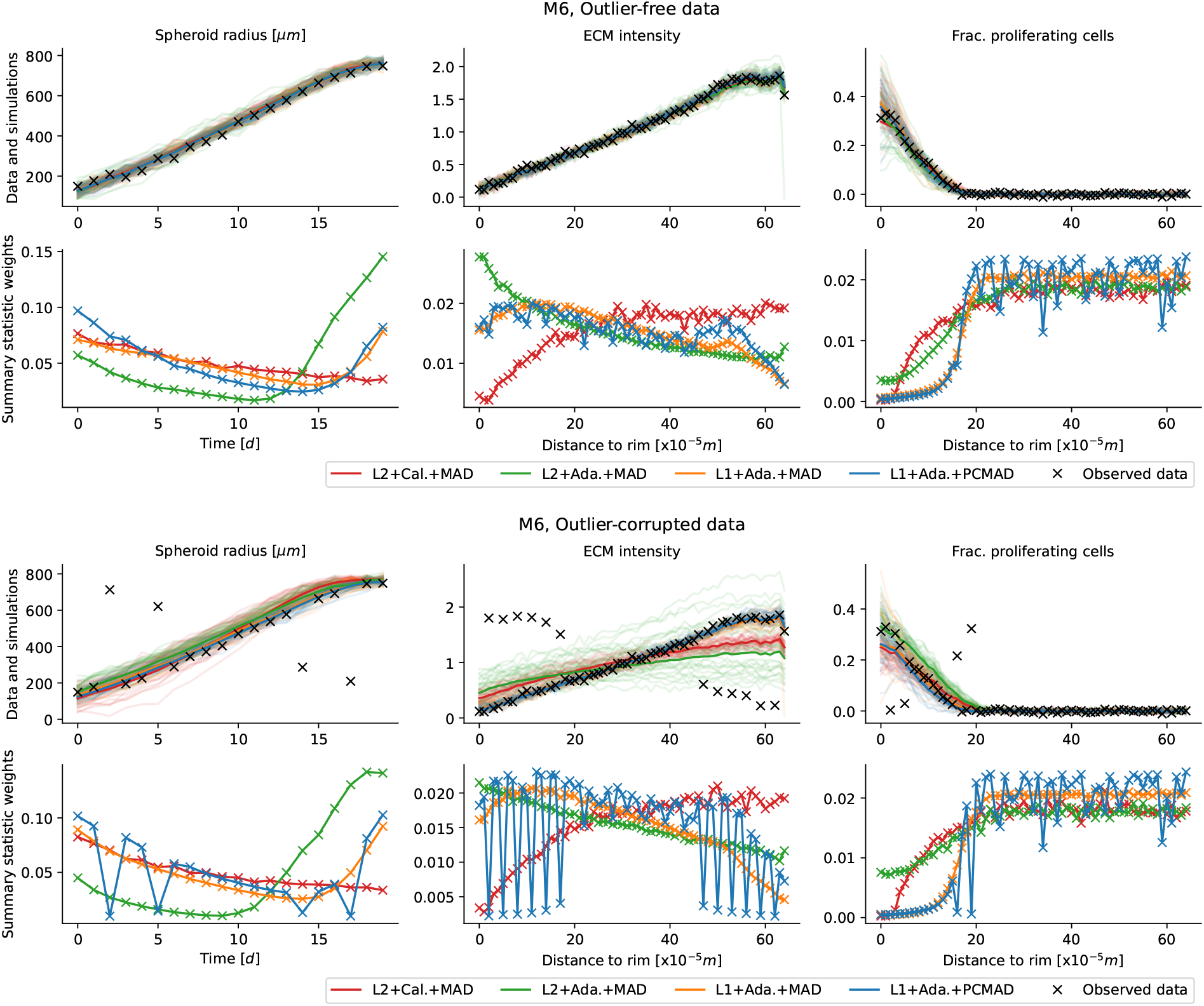
Fits and weights for four distance functions on problem M6 on outlier-free (top) and outlier-corrupted (bottom) data. The respective upper rows show the observed data (black), and, for each distance, 30 accepted simulated data sets (light lines) as well as the sample means (darker lines) from the last ABC-SMC generation. Note that these are accepted simulations, not predictions; for *ε* → 0, the accepted simulations should exactly match the observed (non-outlier) data. The respective lower rows show the corresponding weights assigned to each summary statistic by the four distance functions in the last generation, normalized to sum 1.

On outlier-corrupted data, both L2+Cal.+MAD and L2+Ada.+MAD performed badly, with highly uncertain or unreasonable parameter estimates (Figure 4 bottom), and simulations that visually do not fit to the observed data (Figure 5 bottom). In contrast, L1+Ada.+MAD and L1+Ada.+PCMAD performed considerably better, with parameter estimates closer to those obtained on outlier-free data, although generally with higher uncertainties as expectable given the high fraction of outliers, and better data fits. Overall, L1+Ada.+PCMAD provided the smallest parameter uncertainties on all parameters. Further, the outlier detection reliably identified and corrected for virtually all outliers.

## 4 Discussion

ABC methods enable easily accessible likelihood-free inference for complex stochastic models. However, we have shown in this work how established approaches may be sensitive to outliers in the data. An extension to a popular distance function with iteratively updated scale weights, we have introduced a robust adaptive distance function based on an L1 norm, and further introduced a weighting scheme that allows for the simulation-based online detection and down-weighting of outliers. We evaluated and compared the novel methods on six test problems. This demonstrated firstly that the use of an outlier-insensitive distance function such as L1 considerably improves performance in the presence of outliers, while performing similarly well on outlier-free data. Secondly, the outlier detection and correction generally further improved results, as outliers were correctly identified and sufficiently down-weighted, allowing the analysis to extract information from the relevant data more efficiently. Also against extreme outliers or model misspecification, which severely affected reference methods, did the presented methods prove robust. As especially in high-dimensional complex data sets, e.g. based on imaging techniques commonly used in ABC applications, outliers or highly deviant data points may be expected to exist, we recommend, given our results, the consistent use of robust distance functions in practical applications, unless there are reasons against, e.g. if sensitivity to large deviations is desirable. Further, also the here introduced outlier correction method appears generally advisable, if only for detection, in order to inform early-on about potential problems in data or model.

One possible path of future research would be the evaluation of distances that combine the sensitivity of L2 distances to small deviations with the robustness of L1 distances to large outliers, such as a Huber distance, which however relies on the choice of hyperparameters. Also a comparison to alternative robust distance functions as mentioned in the introduction is beyond the scope of this work, however would definitely be of interest, wherever multiple approaches are applicable.

To ensure convergence, distance weights need to be either frozen after a number of generations, or their eccentricity bounded. Devising reliable strategies of doing so while allowing for sufficient weight adaptation would be practically relevant.

A further promising path becomes apparent in the last application example (Figure 5): One data type, the fraction of proliferating cells, is constant over most of the observed data points. Thus, these summary statistics are hardly informative of the parameters, yet are assigned high weights due to their low variability. Thus, measuring the “informativeness” of summary statistics in combination with scale adaptation and outlier correction might considerably improve the analysis in such cases.

In summary, we have presented a novel robust adaptive distance function with outlier downweighting, and demonstrated its broad applicability. The methods are easy to adopt and have been implemented in an openly accessible tool allowing massive parallelization, facilitating the straightforward use in application projects.

## Acknowledgments

The authors gratefully acknowledge the Gauss Centre for Supercomputing e.V. (https://www.gauss-centre.eu) for funding this project by providing computing time on the GCS Supercomputer JUWELS [Jülich Supercomputing Centre, 2019] at Jülich Supercomputing Centre (JSC).

## Author contributions

Y.S. derived the theoretical foundation and devised the algorithms, wrote the implementation and performed the case study. E.A. assisted with the automated execution of the case studies on high-performance infrastructure. J.H. conceived the research question and provided supervision. All authors discussed the results and conclusions and jointly wrote and approved the final manuscript.

