## Supplementary information for "Robust adaptive distance functions for approximate Bayesian inference on outlier-corrupted data"

#### 1 Test problems

In this section, we give details on all test problems considered in this work, including model description, ground truth parameters, priors, and outlier scenarios.

##### 1.1 Uninformative problem M1

This problem consists of 11 variables, thereof $s_j \sim \mathcal{N}(\theta, 1)$ for $j = 1, \dots, 10$, and $s_{11} \sim \mathcal{N}(5, 0.1^2)$. Ground truth parameter is $\theta = 5$, prior $\theta \sim U[0, 10]$. As outlier we fixed $s_{11} = 7$.

##### 1.2 Replicates problem M2

This problem consists of 10 variables $s_j \sim \mathcal{N}(\theta, 0.2^2)$, with ground truth parameter $\theta = 6$, prior $\theta \sim U[0, 10]$. As outlier we set two variables to $s_j = 0$.

##### 1.3 Conversion problem M3

This problem is based on an ODE model of a simple biological conversion reaction,

$$A \underset{\theta_2}{\overset{\theta_1}{\rightleftharpoons}} B,$$

the analytical solution to which is

$$\begin{pmatrix} A \\ B \end{pmatrix}(t, \theta) = \frac{1}{\theta_1 + \theta_2} \left[ \begin{pmatrix} \theta_2 & \theta_2 \\ \theta_1 & \theta_1 \end{pmatrix} - \begin{pmatrix} -\theta_1 & \theta_2 \\ \theta_1 & -\theta_2 \end{pmatrix} \exp(-(\theta_1 + \theta_2)t) \right] \begin{pmatrix} A_0 \\ B_0 \end{pmatrix}$$

with parameter vector $\theta = (\theta_1, \theta_2)$. We assumed initial concentrations $(A_0, B_0) = (1, 0)$ and only species $B$ to be measured. We assumed to have $n_y = 10$ equidistant measurement points in the interval $[0, 60]$. The measured data were assumed to be noise-corrupted by additive normal noise with a standard deviation of $\sigma = 0.02$, thus we added that random variable to simulation outputs. Both reaction rate coefficients were estimated on log-scale, with independent uniform priors $(\log\theta_1, \log\theta_2) \sim U[-3.5, 1]^{\bigotimes 2}$. Ground truth values were $(\log\theta_1, \log\theta_2) = (-1.5, -1.5)$. This problem is based on and mostly identical to a test problem in Maier et al. [2017].

##### 1.4 GK problem M4

This problem is based on the g-and-k distribution, which is defined via its quantile function

$$q(x) = A + B\left(1 + c\frac{1-\exp(-gz(x))}{1+\exp(-gz(x))}\right)(1+z(x)^2)^k z(x),$$

where $z(x)$ is the quantile function of a standard normal distribution. It does not have a closed-form likelihood function, but can be sampled from by sampling $z \sim \mathcal{N}(0, 1)$ and substituting. We fixed $c = 0.8$ and estimated the remaining parameters. We used as summary statistics seven order statistics at indices $(1250, 2500, ..., 8750)$ out of $10,000$ independent samples. We used independent $U[0, 10]$ priors on the model's four parameters $A, B, g, k$, and ground truth values $(A, B, g, k) = (3, 1, 1.5, 0.5)$. As outlier we considered randomly setting one observable to zero. This problem is based on and mostly identical to a test problem in Prangle [2017].

##### 1.5 Lotka-Volterra problem M5

This problem is a Markov jump process model of a Lotka-Volterra predator-prey population model. The underlying simplified population dynamics of prey $x_1$ and predators $x_2$ are given via

$$\begin{aligned}
(x_1, x_2) &\overset{\theta_1}{\to} (x_1 + 1, x_2) && \text{(prey growth)},\\
(x_1, x_2) &\overset{\theta_2}{\to} (x_1 - 1, x_2 + 1) && \text{(predation)},\\
(x_1, x_2) &\overset{\theta_3}{\to} (x_1, x_2 - 1) && \text{(predator death)}.
\end{aligned}$$

Simulations were performed using Gillespie's direct algorithm [Gillespie, 1977], using the python package `ssa` (`https://pypi.org/project/ssa/`) developed by the authors. We assumed both species to be observed under independent additive normal noise of variance $\exp(2.3)$, at 16 equidistant time-points $t = 2, 4, \ldots, 32$, over a span of roughly four oscillations. The model has 3 rate coefficients, which were estimated on log-scale, with independent $U[-6, 2]$ priors. As ground-truth we considered $(\theta_1, \theta_2, \theta_3) = (1, 0.005, 0.6)$. As outliers we considered inverting the sign of all 6 observables at 3 random time-points. This problem is based on and mostly identical to a test problem in Prangle [2017].

##### 1.6 Tumor problem M6

This problem is a multi-scale agent-based model of tumor spheroid growth on a two-dimensional plane, as introduced in Jagiella et al. [2017]. Single cells are modeled as stochastically interacting agents, coupled to the dynamics of extracellular substances modeled via partial differential equations. The model describes three observables: The spheroid radius over time, and the extra-cellular matrix (ECM) density and the fraction of proliferating cells, at different distances from the rim, observed at a single time point. The model is written in `C++` with python bindings and available at `https://github.com/icb-dcm/tumor2d`. On top of the agent-based model, we assumed independent normal measurement noise of standard deviations 15, 0.04, 0.006 for growth curve, ECM profile, and proliferation profile, respectively. The model possesses seven unknown parameters: Division rate, initial spheroid radius, initial fraction of quiescent cells, division depth, ECM production rate, ECM degradation rate, ECM division threshold. All parameters were estimated on log-scale with default independent uniform priors $U[-3,-1]\times U[1,3]\times U[0,1.2]\times U[-5,0]\times U[-5,0]\times U[-5,0]\times U[-5,0]$. Ground truth values were $(4.17\cdot 10^{-2}, 1.2\cdot 10^{1}, 7.5\cdot 10^{-1}, 10^{2}, 5\cdot 10^{-3}, 8\cdot 10^{-4}, 10^{-2})$. As outliers we considered interchanging 20 of the data points in the observables' dynamic regimes, explicitly every third data point from the back and the front.

### 2 Performance using only the current generation's acceptance criterion

In order to assess the effect of nested acceptance regions on robustness, we performed the same analyses shown in the main manuscript, but basing the acceptance decision only on $d^t(s, s_{\text{obs}}) \leq \varepsilon_t$, instead of checking also previous acceptance criteria. The corresponding plots are shown in this section in Figures S1, S2, S3.

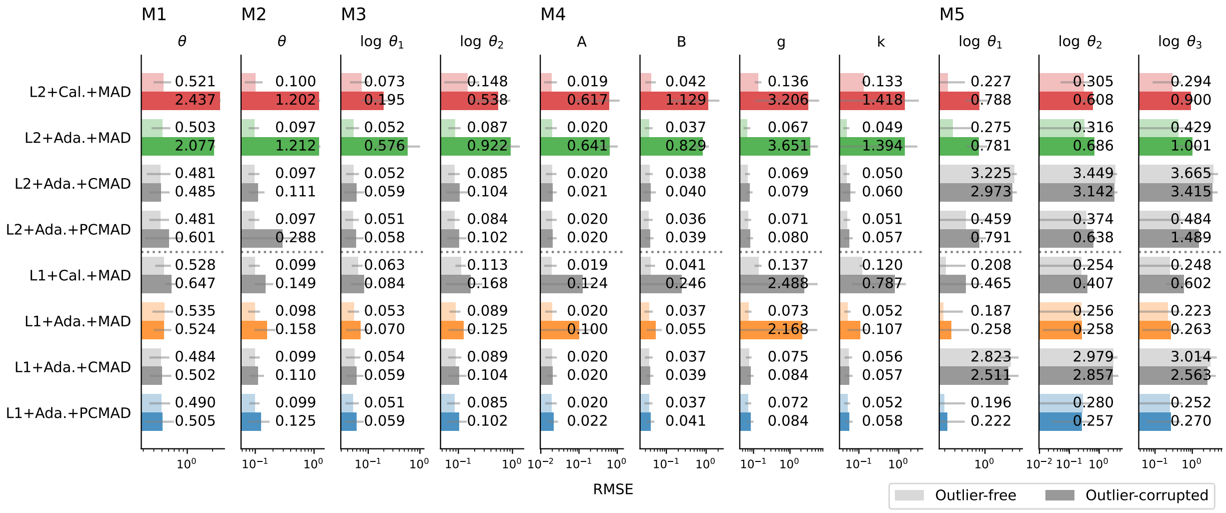


Figure S1: Mean RMSE for the parameters of 5 test models (columns) obtained for 8 distance functions (rows), using L2 or L1 distances, calibrated only in the first ("Cal.") or every ("Ada.") generation, and using MAD, CMAD, and PCMAD for distance weight calculation. Each RMSE is averaged over 20 data sets, grey lines indicate standard deviations. For each distance, the upper, lighter bar is based on outlier-free data, while the lower, darker bar is based on outlier-corrupted data. Distances of interest are colored, alternative distance combinations are shown in grey for reference. Unlike the main manuscript, here previous generations' acceptance criteria were not taken into account.

As can be seen from the comparison of Figure S1 with its pendant in the main manuscript, a structural advantage of either approach cannot be observed especially for the L1 distances. Both perform better in different cases and similarly in most. Slightly better results are generally obtained using previous acceptance criteria. Only on the Lotka-Volterra problem does taking the history into account seem to provide a consistent advantage, indicating that a more conservative choice of weighting with nested acceptance regions may be of advantage on highly flexible problems. Note that on this example, pre-calibrated weights already give comparable results.

For model M6 (Figures S2 and S3), L2+Ada.+MAD performs considerably worse when not using previous acceptance criteria, while the difference is far less pronounced for the L1 distances.

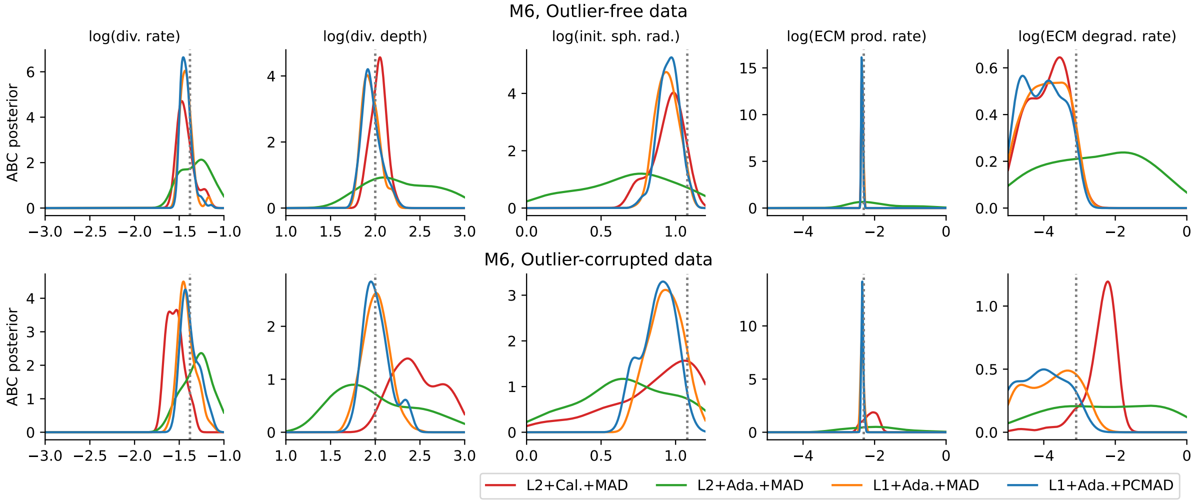


Figure S2: Posterior marginals for the 5 out of the 7 model parameters of model M6 showing interesting dynamics. Top row: Without outliers. Bottom row: With outliers. The x-axis boundaries are the uniform prior boundaries. The parameter values used to simulate the observed data are indicated by grey dotted lines. This is a replicate of the corresponding figure in the main manuscript, not using previous generations' acceptance criteria.

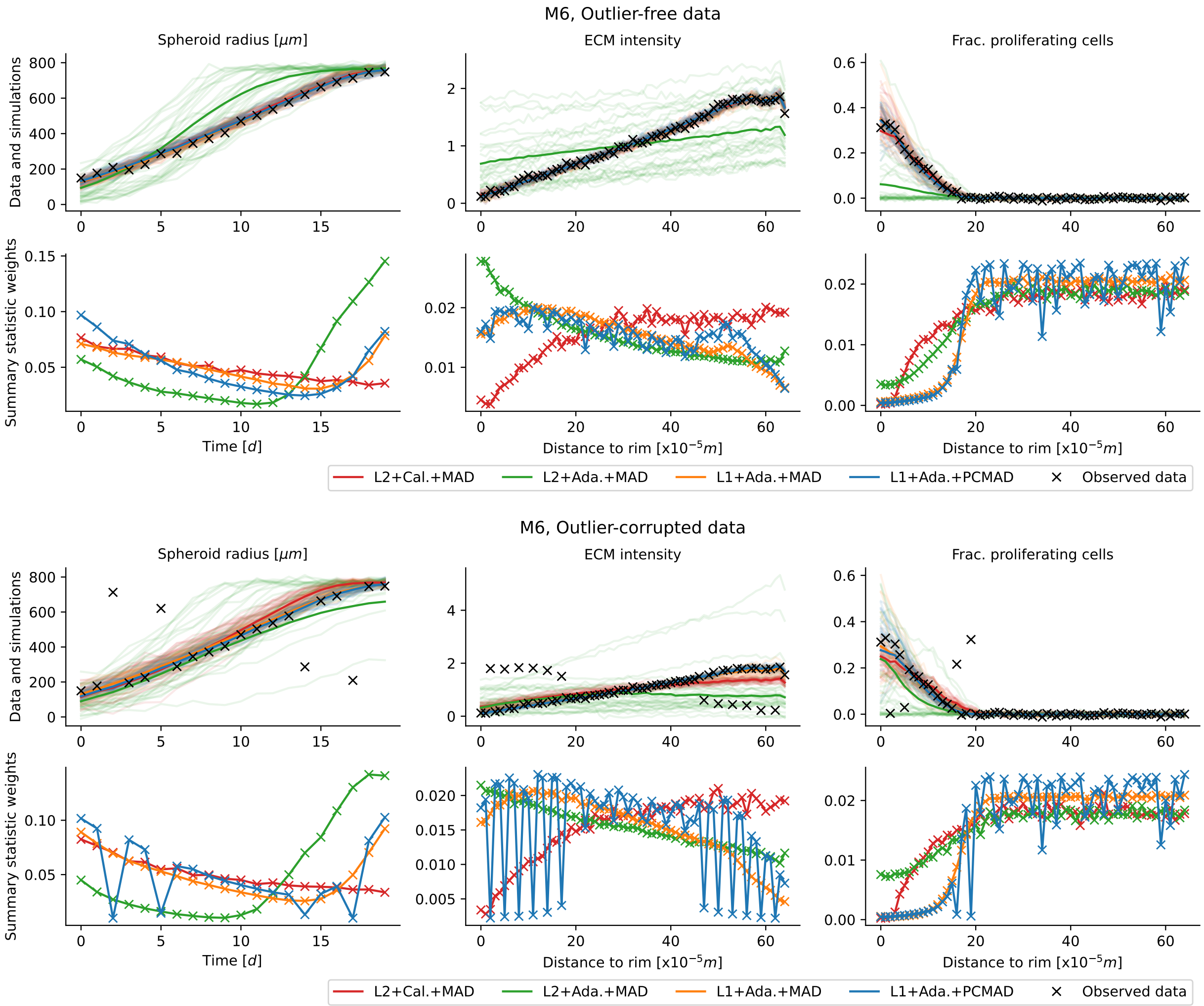


Figure S3: Fits and weights for four distance functions on problem M6 on outlier-free (top) and outlier-corrupted (bottom) data. The respective upper rows show the observed data (black), and, for each distance, 30 accepted simulated data sets (light lines) as well as the sample means (darker lines) from the last ABC-SMC generation. Note that these are accepted simulations, not predictions; for $\varepsilon \to 0$, the accepted simulations should exactly match the observed (non-outlier) data. The respective lower rows show the corresponding weights assigned to each summary statistic by the four distance functions in the last generation, normalized to sum 1. This is a replicate of the corresponding figure in the main manuscript, not using previous generations' acceptance criteria.

### 3 Impact of outlier scale

To study the impact of the scale of deviations, we have run the test problems M1 and M2 with outliers of 1, 5, 25, 125 times the corresponding statistics' standard deviations. Figure S4 summarizes the results.

As expected, larger deviations have a greater influence on L2 distance and thus impact the analysis more severely, leading to larger uncertainties or more biased results. Both here suggested L1-based distances are affected far less.

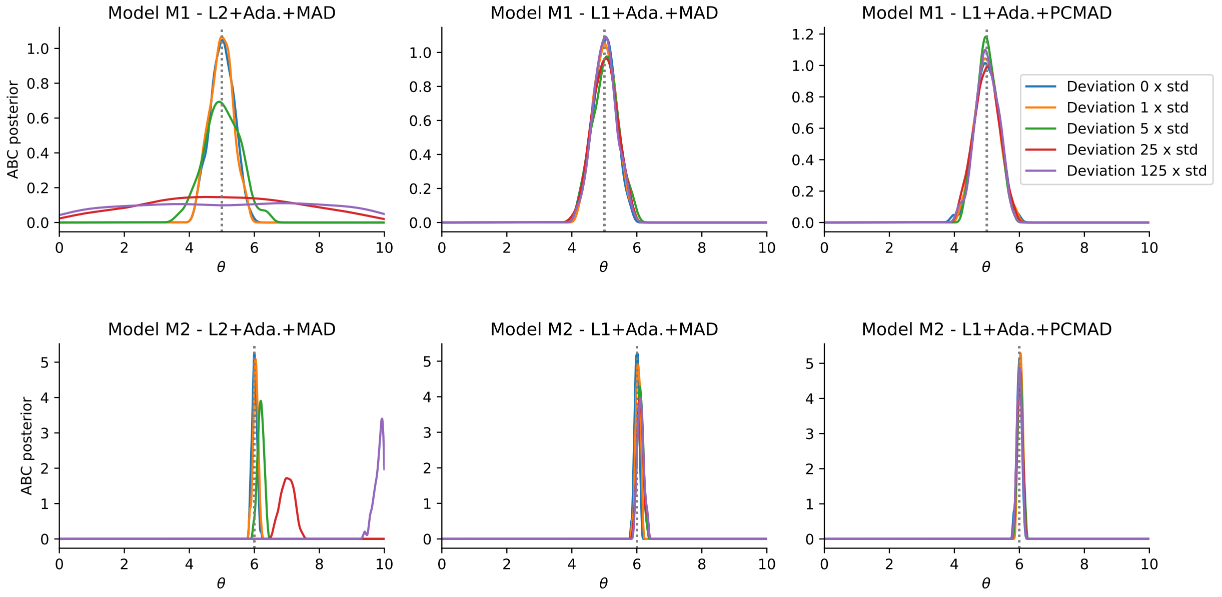


Figure S4: Impact of scale of deviation on the analysis. Top: Test problem M1. Bottom: Test problem M2. Left to right: Adaptive L2 norm with MAD weights, adaptive L1 norm with MAD weights, and adaptive L1 norm with PCMAD weights. Shown are five outlier scales with perfect data at the distribution means, except for the outlier data points which were given deviations of 0, 1, 5, 25, 125 times the standard deviation of the respective summary statistics. True parameters ($\theta = 5$ for M1, $\theta = 6$ for M2) are indicated by grey dotted lines.
